# Optimal sequencing depth for measuring the concentrations of molecular barcodes

**DOI:** 10.1101/2024.06.02.596943

**Authors:** Tommaso Ocari, Emilia A. Zin, Muge Tekinsoy, Timothé Van Meter, Chiara Cammarota, Deniz Dalkara, Takahiro Nemoto, Ulisse Ferrari

## Abstract

In combinatorial genetic engineering experiments, next-generation sequencing (NGS) allows for measuring the concentrations of barcoded or mutated genes within highly diverse libraries. When designing and interpreting these experiments, sequencing depths are thus important parameters to take into account. Service providers follow established guidelines to determine NGS depth depending on the type of experiment, such as RNA sequencing or whole genome sequencing. However, guidelines specifically tailored for measuring barcode concentrations have not yet reached an accepted consensus. To address this issue, we combine the analysis of NGS datasets from barcoded libraries with a mathematical model taking into account the PCR amplification in library preparation. We demonstrate on several datasets that noise in the NGS counts increases with the sequencing depth; consequently, beyond certain limits, deeper sequencing does not improve the precision of measuring barcode concentrations. We propose, as rule of thumb, that the optimal sequencing depth should be about ten times the initial amount of barcoded DNA molecules before any amplification step.

## Introduction

Since its commercialisation in the early 2000s, next generation sequencing (NGS) allows to sequence millions of DNA molecules in parallel [1, 2], revolutionizing nucleic acid research. Since then, NGS cost has constantly dropped [3], and today it represents a powerful and accessible tool for many research investigations. In protein engineering, NGS has radically changed experimental approaches to study the fitness landscape of mutant proteins, as in deep mutational scanning experiments (DMS) [4, 5, 6, 7, 8]. It has also shifted the paradigm of designing new proteins with augmented functions in systematic evolution of ligands by exponential enrichment (SELEX) [9, 10, 11, 12, 13] or in directed evolution (DE) [14, 15, 16, 17, 18, 19, 20]. In these experiments, highly diverse gene libraries are initially prepared, where each gene is tagged with a unique molecular barcode or a distinctive mutation. Then, the fitness of each variant is assessed by measuring the relative concentration of each barcode using NGS (Fig. 1a), before and after one or multiple screening rounds. For example, in DMS experiments, a large library with thousands, if not millions of mutants undergo a screening [5], and NGS before and after a selection experiment distinguishes between beneficial and detrimental mutations, as those that increase or decrease their concentration [6, 7].

**Fig. 1:**
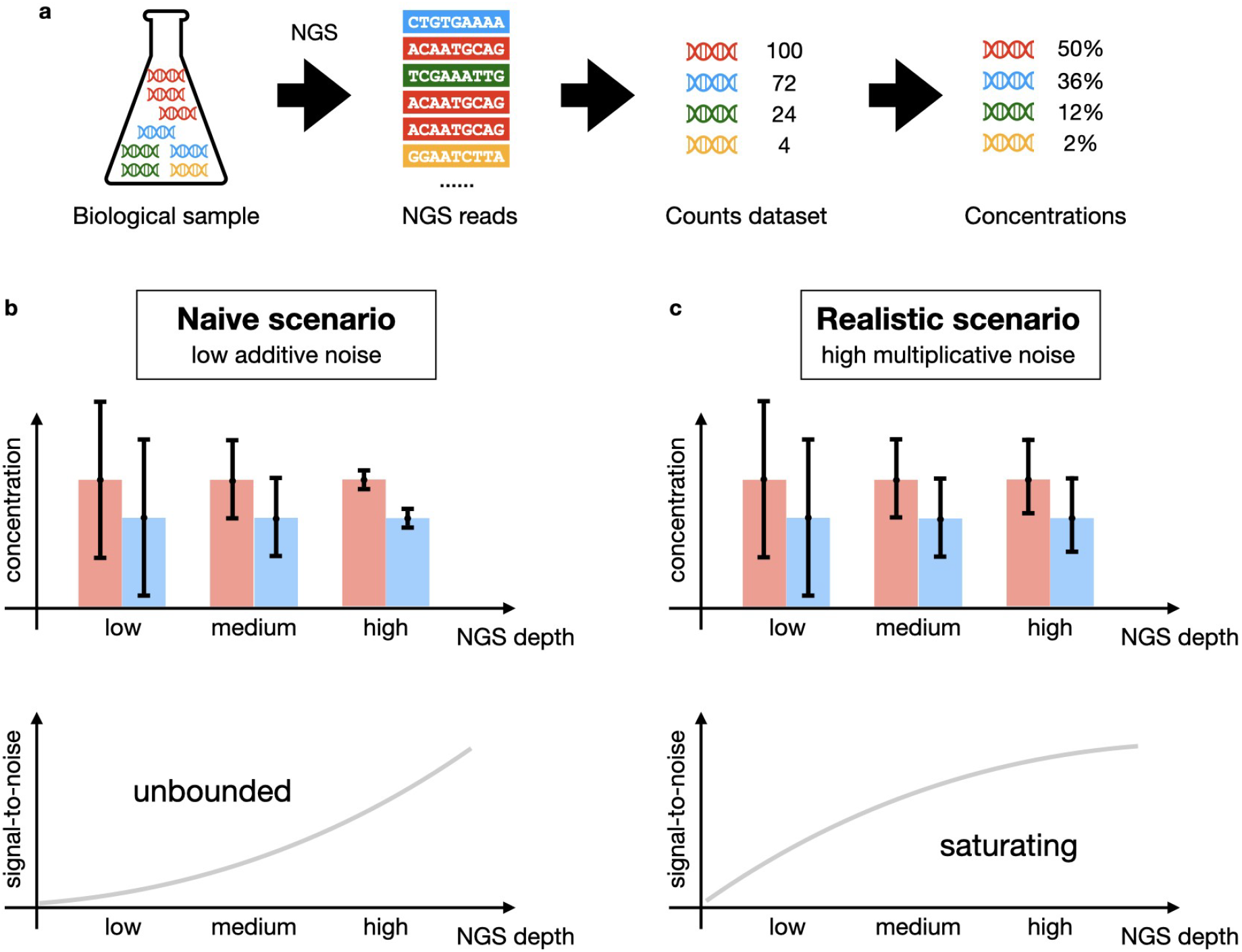
Statistical errors in sequencing analysis. (a) NGS is often used to compute concentrations of different species or molecular barcodes in highly diverse DNA libraries. (b) Naïve scenario underestimates noise in NGS counts, expects that statistical errors can be reduced by enlarging NGS depths, and therefore the signal-to-noise reatio be increased indefinitely. (c) In the realistic scenario evinced in this work, errors do not always shrink with deeper NGS, and signal-to-noise saturate after a certain depth.

Being able to precisely measure the concentrations of the library content is therefore crucial. For this reason, the number of NGS reads (NGS depth) is an important parameter to both the design and interpretation of the experiments. In other applications of NGS, such as whole genome sequencing or RNA sequencing, the optimal NGS depth has been historically discussed intensively in terms of coverage [21], and guidelines have been established [22, 23]. Yet, for the measurements of barcode concentrations, such guidelines are not yet available and the NGS depth is often pushed to the limit, resulting in non-negligible costs for the laboratories. Here, we point out that this is not beneficial for most cases. Increasing the NGS depth does not always reduce the impact of random sampling and therefore does not shrink the statistical errors (*naïve scenario*, Fig. 1b).

A standard workflow to prepare a DNA-sample library for NGS includes PCR amplifications. Even highly reliable PCRs introduce errors during DNA replication, and they are of different nature. During a PCR amplification, some DNA molecules might not get replicated, or some wrong nucleotides may be inserted randomly [24]. If this happens at the beginning of the amplification protocol then the error will be amplified as well [25, 26, 27, 28, 29]. In this work we first analyze the structure of these amplification biases in NGS datasets, to then develop a mathematical model that accounts for it. Built on previous works [30, 31, 32, 33, 34, 35], our model accounts for how PCR mistakes impact the whole NGS process. This analysis showcases that, by increasing the NGS depth, both signal and noise increase in the estimation of concentration in highly diverse libraries. As a consequence, beyond a certain depth – approximately ten times the initial amount of molecules of barcoded DNA (from now on simply called initial DNA amount) – deeper NGS does not increase data quality (*realistic scenario*, Fig. 1c), and this sets an optimal number of reads that need to be chosen to obtain the most from the data, without wasting resources.

## Materials and Methods

### Dataset characteristics

In this work we considered nine barcode libraries with different characteristics (Table 1). Three datasets (*AAV dataset 1, 2, 3*) used random 7-mer peptides as barcodes, inserted between amino acid 587 and 588 of the cap2 gene in the adeno-associated viruses serotype 2 (AAV2) plasmid. *AAV dataset 1* used random 7-mer peptides composed of six VNN codons (N stands for any base, V stands for any base but Thymine) and one VNG codon (G stands for Guanine only) [29], where the distribution of amino acids is biased towards a higher abundance of lysine (Fig S1). *AAV dataset 2* used random 7-mer peptides of seven NNK codons (K stands for Guanine or Thymine base), where all admitted nucleotides are designed to have the similar occurrence. This led to relatively flat amino-acid distributions [36] (Fig S1). *AAV dataset 3* is our in-house dataset prepared using Twist Bioscience^®^ as described below, and resulted in the most homogeneous occurrence of all amino-acids at all of the 7 positions (Fig S1) among the three datasets. NGS was used to sequence the barcoded regions of these libraries, where the number of unique sequences and NGS depths are summarised in Table 1. Other two datasets (*hYAP65 WW rep 1, 2*) are made of barcodes of the mutated WW domain of the hYAP65 protein (99 nucleotides of variable region). These datasets were generated through error-prone PCR, so their counts decrease with the number of mutations from the wild-type sequence. The last four datasets (*scRNAseq-UMI 1, 2, 3, 4*) used as barcodes the unique molecular identifiers (UMI) of publicly available single-cell-RNA-sequencing datasets from 10X Genomics database. We observed that nucleotide distributions are relatively uniform, though GC-ratio is smaller than 50% in all of scRNAseq-UMI datasets (Fig S1). Note that all nine datasets are made of oligos (barcodes). The different names are related to the template where the barcodes are inserted. However the templates do not influence the analysis which follows here.

**Table 1.** Summary of datasets. Nucleo-flat (amino-flat) means that the nucleotides (amino-acids) showed similar frequencies across different position. The first dataset was biased toward an high adenine content, resulting in an highly heterogeneous amino-acid content.

| Dataset | Length | Unique sequences | NGS depth | Probability distribution |
| --- | --- | --- | --- | --- |
| AAV dataset 1[29] | 7 amino acids | 1,252,032 | 13,653,491 | biased |
| AAV dataset 2[36] | 7 amino acids | 3,071,451 | 12,195,473 | nucleo-flat |
| AAV dataset 3 | 7 amino acids | 98,468,090 | 240,537,294 | amino-flat |
| hYAP65 WW rep 1[38] | 99 nucleotides | 1,336,842 | 3,588,061 | error-prone PCR |
| hYAP65 WW rep 2[38] | 99 nucleotides | 1,615,935 | 4,931,094 | error-prone PCR |
| scRNAseq-UMI 1[39] | 16 nucleotides | 14,512,602 | 75,820,168 | nucleo-flat |
| scRNAseq-UMI 2[40] | 16 nucleotides | 16,020,053 | 173,326,275 | nucleo-flat |
| scRNAseq-UMI 3[41] | 16 nucleotides | 16,429,852 | 312,213,184 | nucleo-flat |
| scRNAseq-UMI 4[42] | 16 nucleotides | 14,974,774 | 182,330,818 | nucleo-flat |

For AAV datasets, cutadapt [37] was used to extract sequences that correspond to random 7-mer peptides from FASTQ files. The average error rates are estimated for each sequence from Q score and those that have more than 0.1 error rates are removed for further analysis. Translating nucleotide sequences into amino acid sequences, 7 amino acids are obtained as barcodes. In this step, sequences containing stop codon and amino acids that cannot be formed due to the codon design are removed. For the hYAM65 datasets forward and reverse reads were compared and if the two reads match the entire sequence was kept in the dataset. For scRNAseq-UMI datasets, BAM files are downloaded from the 10X Genomics database, and UMI with UR tags consisting of 16 nucleotides are extracted. These UMIs are used as barcodes.

### Amino-flat library preparation

We ordered 50 *µ*g of a AAV plasmid library from Twist Bioscience^®^ that used silicon-based DNA synthesis platform. Ten amino acids were inserted at position 587/588 in the AAV2 Cap sequence, where three of them were the AA/A linkers and seven of them, flanked by AA/A, were random peptides consisting of a diversity of 10^8^ to 10^9^ variants. The random peptides were inserted in the way that each amino acid occurred at the same probability in each position. We requested that the restriction sites for restriction enzymes HindIII, NotI and AscI be kept in the original AAV2 Rep/Cap plasmid sequence to permit future cloning.

10 ng of the 7mer library was PCR amplified using primers 5 ‘-ATCAGGACAACCAATCCCGTGGCTA-3’ and 5’-TGTCCT GCCAGACCATGCCTG-3’ with Takara Bio PrimeSTAR GXL high-fidelity polymerases. After the PCR amplification, reactions were cleaned-up with a Macherey-Nagel Gel Extraction and PCR Clean-Up kit. Samples were submitted to Plateforme GENOM’IC at Institut Cochin for sequencing. They were sequenced with Illumina NextSeq 2000, P3 1*200 cycles, at 200 million reads per sample.

### A position-independent model

To analyze the data AAV and scRNAseq, we use a position-independent or position weight model [43], which serves as a proxy for the original concentrations of each barcode in the samples. In this model, we assume that each barcode sequence *s* = *s*^1^*s*^2^ … *s*^*L*^, where *L* is the length of the barcode, is generated from the following probability: 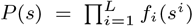, where *f*_*i*_(*s*^*i*^) is the frequency of *s*^*i*^ at the position *i* estimated from each dataset. To analyze the AAV datasets, we use amino acids sequence: *s*^*i*^ ∈ (20 amino acids) with *L* = 7, whereas for the scRNAseq-UMI datasets, we use nucleotides sequence: *s*^*i*^ = *T, A, G, C* with *L* = 16.

### A mutation model

To analyze the data of error-prone PCR we use a position-independent mutation model. In this model we assume that each barcode sequence is uniquely described by the number and the type of mutations from the wild-type barcode 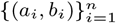 where *n* is the number of total mutations and *a*_*i*_ is the nucleotide of the wild-type which is mutated into the nucleotide *b*_*i*_. The parameters of the models are the vector *p*(*n*) which is the probability of having exactly *n* mutations and the matrix *M* (*a, b*) which gives the probability of replacing a nucleotide *a* = *A, C, G, T* with the nucleotide *b ≠ a*. The probability of a barcode to be in the dataset is indeed 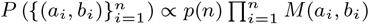.

### Calculation of the noise extent (Fano factor) from data

Either using the position-independent model or the mutation model, the expected count *λ* (*s*) of a barcode sequence *s* is computed by multiplying the sequence probability *P* (*s*) by the sequencing depth *D*: *λ* (*s*) = *DP* (*s*). Note that the expected count can be estimated for all possible barcode sequences, including those that have zero observed counts in the dataset. Next, barcode sequences are binned based on their expected count *λ*(*s*) in *N*_*b*_ bins, such that each bin has the same number of barcode sequences with nonzero observed counts. Note that the binning would have been infeasible on the observed counts given their discrete nature. The average value and variance of the observed counts in each bin are denoted by ⟨*c*⟩_bin_ and *σ*^2^_*c*,bin_. Similarly, those of the expected counts *λ*(*s*) in each bin are denoted by ⟨*λ*⟩_bin_ and *σ*^2^_*λ*,bin_. Using these quantities, the extent of noise in the data is computed as the ratio between variance and mean, commonly referred to as the Fano factor (F.F.). The Fano factor is a quantity related to the amount of dispersion we have in the data, since a higher F.F. corresponds to a larger variance given the same mean. Here, taking into account the variance of *λ* in each bin, we estimate it as

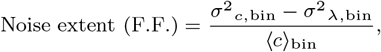

Note that the difference at the numerator is due to the fact that the expected count *λ* is also a random variable in the bin, so we needed to subtract the variance of this variable. Nevertheless, this is a minor correction, which is negligible for most of the bins. Note that, in *AAV dataset 3* and *scRNAseq-UMI 1, 2, 3 and 4, P* (*s*) is almost uniform because all amino acids (AAV dataset 3) or all nucleotides (scRNAseq-UMI 1, 2, 3, 4) have a similar probability (Fig. S1). Therefore, we set *N*_*b*_ = 1 in these cases.

### Mathematical model of the amplification process

The PCR amplification process is numerically simulated by computing the average number of DNA molecules along the cycles. The model comprises the following three steps:

1. An amount *N*_0_ of the DNA molecules is taken from the initial sample. Because this mimics pipetting a part of the sample, we assumed that this random selection is not correlated, and consequently we modelled it as Poisson sampling.
2. The sample undergoes PCR amplification processes, where each DNA molecule in the sample is replicated with a probability *p*_*n*_ *<* 1 in each round *n*. Errors that change the sequences are not taken into account in this framework. We modelled this process as a series of binomial processes.
3. Finally, *D* random molecules are sequenced using NGS. We modelled it again as a Poisson process.

Note that we use the following way of computing collective quantities at each amplification step without running the complete simulation of the amplification process, which is computationally heavy. We will focus on the case where *p*_*n*_ = (1 + *N*_*n*_*/K*_MM_)^−1^ where *N*_*n*_ is the number of DNA molecules at the *n*-th step, and *K*_MM_ is called Michaelis-Menten constant. The relation between the noise extent (F.F.), the NGS depth *D* and the initial DNA amount *N*_0_ is computed as F.F. = 1 + *a*_*n*_*D/N*_0_ (see Results, Eq. 1), where *a*_*n*_ depends *a priori* on the number of amplification rounds *n*. The method takes in input the initial reduced DNA amount *N*_0_*/K*_MM_ and it outputs, for each of 100 rounds of amplification, the replication probability *p*_*n*_, the reduced DNA amount *N*_*n*_*/K*_MM_ and the values of the function *a* of equation 1, here labeled as *a*_*n*_. It proceeds in an iterative way:

- The reduced DNA amount is initialised to *N*_0_*/K*_MM_.
- The function *a*_*n*_ is initialised to the unity *a*_0_ = 1.
- The replication probability is computed iteratively as *p*_*n*_ = (1 + *N*_*n*−1_*/K*_MM_)^−1^.
- The function *a*_*n*_ is computed iteratively as *a*_*n*_ = 1 + (1 + *p*_*n*_) · (*a*_*n*−1_ − 1) + [*p*_*n*_(1 − *p*_*n*_)*/*(1 + *p*_*n*_)] · [(*N*_0_*/K*_MM_)*/*(*N*_*n*_*/K*_MM_)].
- The reduced DNA amount is updated iteratively as (*N*_*n*+1_*/K*_MM_) = (1 + *p*_*n*_) · (*N*_*n*_*/K*_MM_).

### Dataset resolution and optimal depth

We start defining the signal-to-noise (snr) ratio as the mean expected count of each variant divided by the square root of its variance: 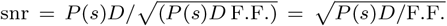, where *P* (*s*) is the probability (concentration) of the variant *s, D* is the sequencing depth and F.F. is the noise extent (Fano Factor). Note that snr is a variant dependent quantity, but the dependence is only via the term 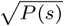. By normalising with respect to it, we can define a resolution as 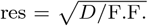 Using our formula for the noise extent (Eq. 1) with *a* = 1, the resolution can be computed as 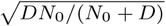. We then define the exploited resolution as the ratio between the resolution and its large *D* limit 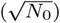 to obtain 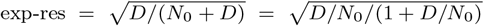. The exploited resolution varies between 0 and 1, depending only on the ratio *D/N*_0_. The optimal depth is computed as the depth *D* for which the exploited resolution reaches 95%. Alternatively, we could have defined an exploited snr, by dividing the snr by its large *D* limit, obtaining 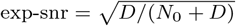, which is equal to the exploited resolution.

### Inference of the number of DNA molecules before amplification

We used Eq. 1 of the Result to estimate the number of initial DNA molecules (*N*_0_) from the sequencing depth (*D*) and the noise extent (F.F.). In the case of replication probability logistically dependent on the amplification rounds, it is possible to approximate *a*_*n*_ ∼ 1. Given noise extent (F.F.) and NGS depth *D*, the DNA amount before amplification *N*_0_ is estimated as *D/*(F.F. − 1).

## Results

### Mean and variance of NGS counts are proportional

We started our analysis by focusing on three barcode datasets where a simple machine learning model provides accurate predictions of the concentrations, and compared them with empirical estimations from NGS counts. We first analyzed *AAV dataset 1* and *AAV dataset 2*, whose NGS depths were approximately 13.6 and 12.2 million reads, resulting in 1.2 and 3.0 millions different observed sequences (Table 1). For each dataset, we inferred a position-independent model (Material and methods), and computed the predicted count for all possible sequences, which amounts to 0.18 and 1.28 billion for *AAV dataset 1* and *AAV dataset 2*, respectively. In order to verify the accuracy of model predictions, we compared the measured counts with those predicted by the model (Fig. 2b). Strong positive correlations were observed in sequences with the empirical count higher than 100. In contrast, sequences with lower empirical counts showed no clear correlations, likely due to their inherently noisy nature. We then sorted the sequences into 200 bins based on their expected counts, grouping those with similar values together (Material and methods). For each bin, the mean of the predicted counts was very close to that of the empirical counts (Fig 2c), showcasing the accuracy of the model predictions. The count variances within each bin, on the other hand, were approximately proportional to the corresponding count means (Fig 2d). We estimated the noise extent of the counts as the Fano Factor (F.F., Material and methods), which corresponds to the variance divided by the mean. The F.F. serves as a measure of noise, with higher values indicating higher variance for a given mean. The noise extent has been computed in each bin, and found that it was significantly larger than 1, indicating that the counts were overdispersed with respect to Poisson noise. The deviation from the proportionality law shown at very high count mean (Fig 2d) is due to the approximations of the model [44] and can at least be decreased by using a more complex model (Fig. S3).

**Fig. 2:**
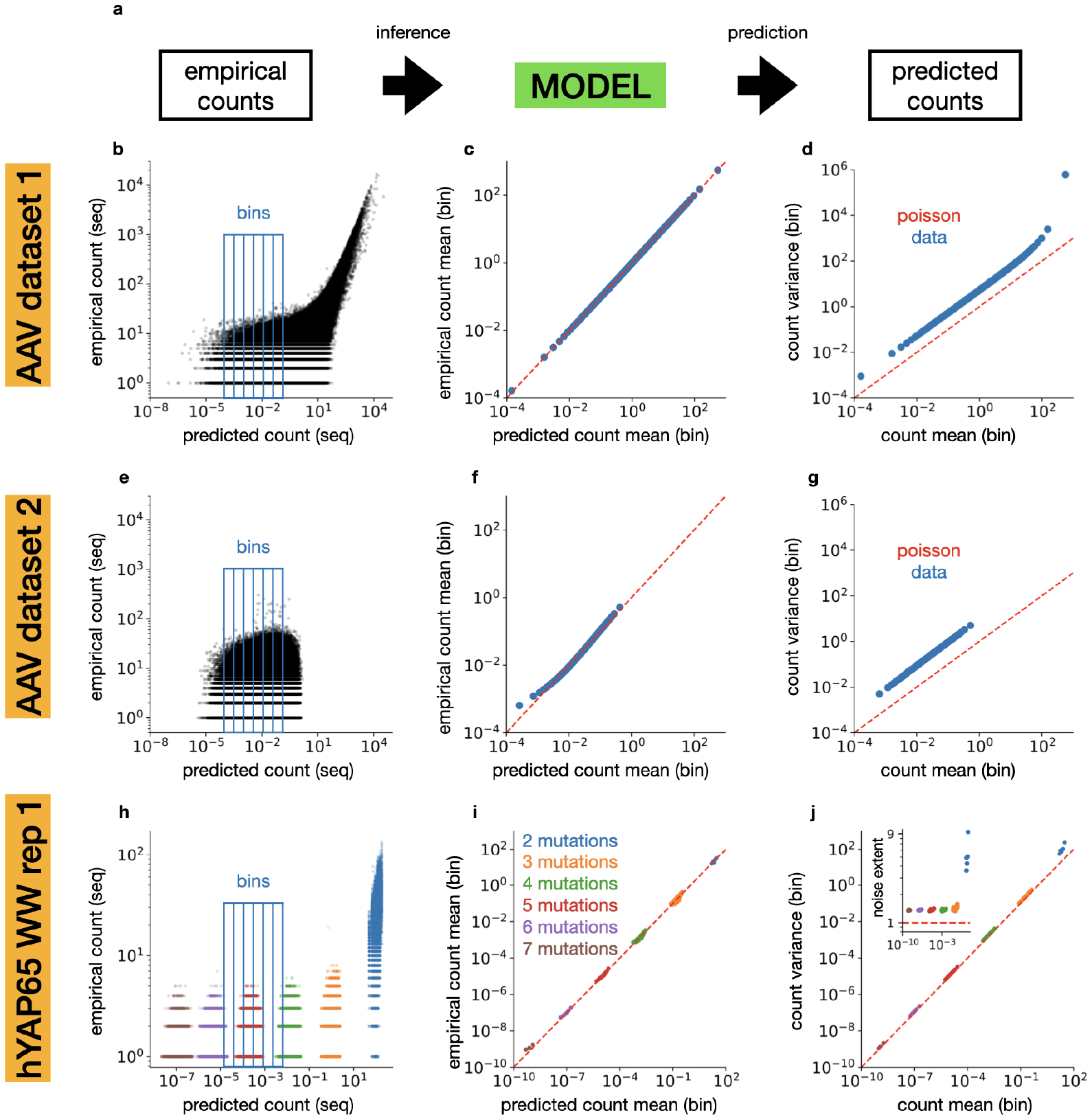
Mean and variance of the counts are proportional in NGS datasets. (a) Starting from the observed counts of molecular barcodes in each NGS dataset we fitted a statistical model to predict the expected count of each sequence. (b) *AAV dataset 1*. Empirical counts plotted against the expected counts for each sequence are shown (black points). Examples of bins grouping sequences with similar predicted counts are also shown as blue boxes. (c) *AAV dataset 1*. Empirical counts and predicted counts were averaged over each bin and are plotted against each other (blue dots). These points align along the equality line (red). (d) *AAV dataset 1*. Variance of the counts against the expected count (where both are averaged over each bin) is displayed by blue dots. In Poisson noise, count variance and expected count are equal to each other (red dashed line). The blue dots align to a line parallel to, yet above, the equality line, thus indicating overdispersion. (e), (f) and (g) are the same as (b), (c) and (d), but use *AAV dataset 2*. (h) same as (b) but use *hYAP65 rep 1* and its corresponding mutation model. (i) same as (f) but use *hYAP65 rep 1* and its corresponding mutation model. Different colors correspond to different number of mutations from the wild-type sequences. (j) same as (g) but use *hYAP65 rep 1* and its corresponding mutation model. Different colors correspond to different number of mutations from the wild-type sequences. In the insight the F.F. (noise extent) for each bin plotted against the mean of the empirical counts.

The analysis of the *AAV dataset 2* provided similar results. Because empirical counts are lower, we did not observe clear correlations between the empirical and predicted counts (Fig. 2e). However, after sorting the sequences into 80 bins based on their expected counts, model predictions of the mean count were well aligned with the empirical values (Fig. 2f). Noise extent (F.F.) was also significantly larger than 1 (Fig. 2g), indicating that the count data were overdispersed.

We finally perfomed the same analysis on the dataset *hYAP65 WW rep 1*. To compute the predicted counts we inferred a mutation model based on the number and the type of mutations from the wild-type sequence (Fig. 2h, Material and methods). Note that the predicted counts are sharply clustered by the number of mutations from the wild-type sequence (different color of Fig. 2h). The noise extent (F.F.) resulted only slightly higher than 1 (Fig. 2j and inset). Similar results came from the analysis of the second replicate (*hYAP65 WW rep 2*, Fig. S2).

### Mathematical model and simulations reproduce the mean-variance proportionality observed in data

In order to increase our understanding of the empirical results found in the previous section, we developed a three-step mathematical model (Material and methods) that mimics the NGS protocol including the PCR amplification process, as detailed in Fig. 3a.

**Fig. 3:**
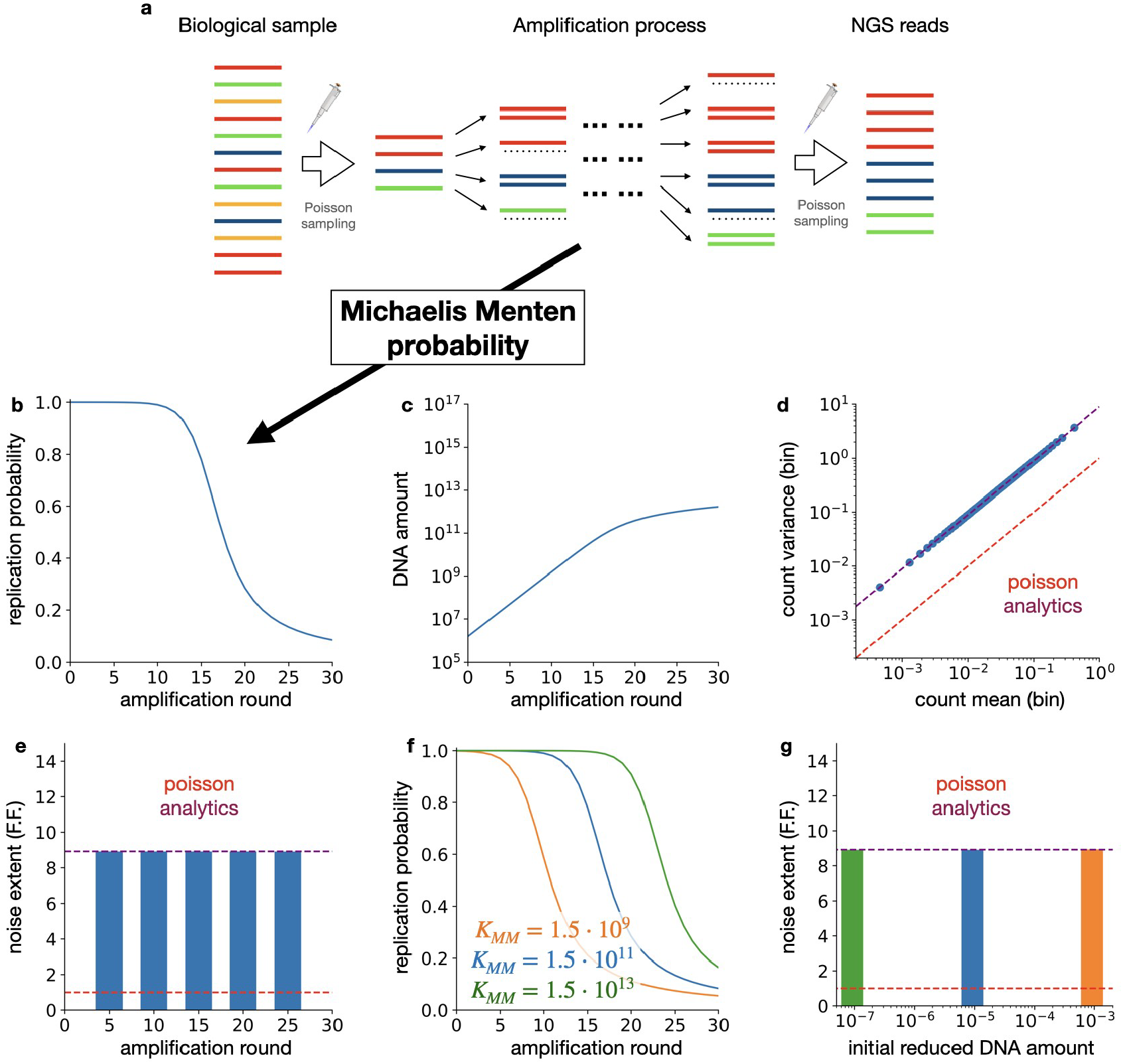
Mathematical model and simulations reproduce the mean-variance proportionality observed in data. (a) The three steps of the mathematical model describing the NGS procedure including PCR amplifications: a poisson sampling is followed by multiple rounds of a binomial process and concluded by another poisson sampling. (b) The replication probability against the number of rounds when the MM constant *K*_MM_ is 1.5 · 10^11^. (c) The total number of DNA barcodes for each round of amplification was simulated, starting from approximately 10^6^ DNA molecules at the 0th round. (d) The count variance against the count mean for each bin of a simulated dataset (blue points) shows overdispersion when compared with Poisson noise (dashed red line) and agrees with the analytical prediction (purple dashed line). (e) Noise extent (F.F.) against the number of amplification rounds confirms its independence on the amplification process. (f) The replication probability against the number of rounds for three different MM constant (1.5 · 10^9^ in orange, 1.5 · 10^11^ in blue, 1.5 · 10^13^ in green). (g) The noise extent (F.F.) is plotted against the initial reduced DNA amount (*N*_0_*/K*_MM_) for the three different MM constants (as shown in the panel f). It suggests the negligible impact of the MM constant in this realistic range..

We first analyzed the model under the simplified assumption of a constant replication probability *p*_*n*_ = *p* across the amplification rounds. Using the theory of Galton-Watson process [45], we computed the noise extent analytically (Fig. S4 and Supplementary mathematics):

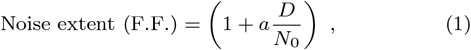

where *D* is the NGS depth, *N*_0_ is the initial number of DNA molecules (from now on also called initial DNA amount (copy number)) before the PCR and here *a* is a constant. This simple mathematical model recovered the proportionality relationship between the mean and variance of the NGS count that we observed in experimental data (Fig. 2). Interestingly, here *a* ≈ 2*/*(1 + *p*) and the noise extent (F.F.) turned out to be independent from the number of amplification rounds. This observation suggests that reducing the number of amplification (such as PCR) rounds does not reduce the noise extent of the final NGS reads (see Discussion).

In order to verify that our analytical results were robust, we relaxed the constant *p* assumption by accounting for the presence of limiting replication factors, such as the quantity of polymerases or primers in PCR. Inspired by previous enzymatic descriptions of PCR process [30, 31, 32], we adopted a replication probability following a Michaelis-Menten (MM) discretized equation (Fig. 3b): *p*_*n*_ = (1 + *N*_*n*_*/K*_MM_)^−1^, where *N*_*n*_ is the total number of DNA molecules present at the beginning of the *n*-th amplification step (DNA amount) and *K*_MM_ is a parameter linked to the limiting factors, known in the literature as the Michaelis-Menten (MM) constant.

We derived an analytical expression for the noise extent in the case of a general replication probability (Supplementary mathematics), and found that it has the same analytical form as before (Eq. 1). Numerical evaluation of the analytical solution with MM replication probability indicates that *a* ≈ 1 holds for a wide range of realistic cases, and in particular for both *K*_MM_ much smaller or much larger than *N*_0_ (Fig. S5). The latter is consistent with the limiting case without replication, for which we obtained *a* = 1 exactly. In the first case, instead, the sample undergoes an initial noiseless replication (*p* ≈ 1) that increases the initial available DNA and therefore shrinks any variability.

To verify our analytical results we conducted numerical simulations of our mathematical model. We initiated the process with a synthetic heterogeneous library containing 20^7^ different sequences, with concentrations given by the inferred probabilities from *AAV dataset 2* (Fig. 2e, f and g). We sampled approximately *N*_0_ ≈ 1.5 · 10^6^ DNA molecules, and we computed the replication probability and the DNA amount for each round up to 30 amplification rounds (Fig. 3b and c). We chose *K*_MM_ ≈ 1.5 · 10^11^ for the MM constant, in order to have an initial *reduced* DNA amount of *N*_0_*/K*_MM_ ≈ 10^−5^. This value of the MM constant lies in the range of realistic PCR, which has been studied to be in between 10^6^ and 10^15^ [46]. Finally, the library was sequenced by sampling the counts with a NGS depth of *D* ≈ 1.4 · 10^7^ reads. The resulting synthetic NGS data were binned over their expected counts - as done in the previous section on real data. We observed an overdispersed behavior with a count variance proportional to the mean, in agreement with analytical predictions (Fig. 3d).

To validate the independence of the noise extent (F.F.) from the amplification processes, we ran five simulations with the same parameters but varying the number of amplification rounds (5, 10, 15, 20 and 25), we computed the noise extent (F.F.) for each case (Materials and methods) and we found similar values for each simulation (Fig. 3e), according to our analytical results.

Finally, to better understand the impact of the MM constant *K*_MM_, we ran three simulations with three different values (*K*_MM_ ≈ 1.5 · 10^9^, 1.5 · 10^11^ and 1.5 · 10^13^). As expected, increasing the constant only shifted the behavior of the replication probability (Fig. 3f). Noise extent was instead constant, in agreement with our prediction (Eq. 1, Fig. 3g).

### Optimal NGS depth

Our mathematical model suggests that noise extent (F.F.) is approximately proportional to the NGS depth (Eq. 1). To validate this observation, we generated eight synthetic datasets by subsequently halving the NGS depth of *AAV dataset 2*. Computing the noise extent after binning (Materials and Methods), we observed a linear relationship between F.F. and the NGS depth as predicted by the mathematical model (Fig. 4a). To further confirm this result, we ran nine simulations of our mathematical model, and analyzed the data with the same procedure. Simulations confirmed the expected linear behaviour and reproduced the empirical analysis (Fig. 4b).

**Fig. 4:**
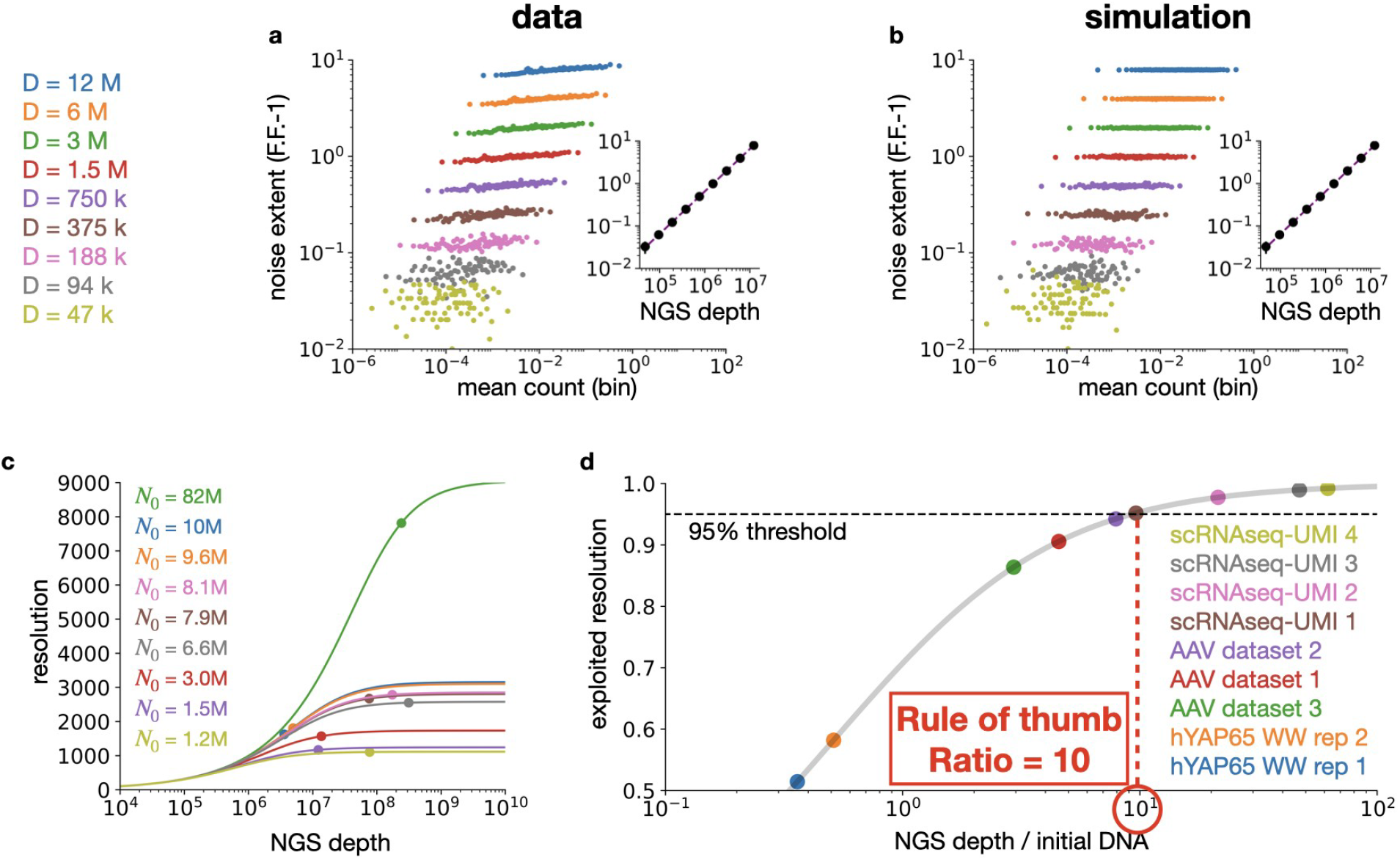
Which is the optimal NGS depth? (a) Eight synthetic datasets were generated from *AAV dataset 2* by subsequently halving the NGS depth. Noise extent (F.F. − 1) is plotted against the average expected counts per bin for each dataset (different colours). Inset: the noise extent (F.F. − 1) against the NGS depth aligns with the analytical predictions (purple dashed line). (b) The same figure as in (a), but obtained from simulating our mathematical model. The parameters for the simulation were tuned to reproduce *AAV dataset 2* (see text for details). (c) Resolution against the NGS depth for nine different initial DNA amounts (*N*_0_), chosen to match the nine datasets of Table 1. Color code is the same as in panel (d). (d) Exploited resolution against the ratio between the NGS depth and the initial DNA (grey curve). The values for the nine datasets (coloured points) are added to compare their depth with the optimal one.

The observation that the noise extent increases with the NGS depth implies that increasing the latter might not always improve data quality, particularly the ability to measure the concentration of DNA barcodes. In order to quantify this ability, we estimated the resolution of a given NGS dataset, as the rescaled signal-to-noise ratio between expected count and its variability (Materials and Method, and Fig.4c). The resolution depends on the two quantities that most affect noise extent, namely the initial DNA copy number and the sequencing depth (respectively, *N*_0_ and *D* in Eq.1). As expected, increasing the amount of initial DNA always increases resolution, or at worst it does not affect it for very low sequencing depth. Interestingly, for a fixed initial amount of DNA, increasing sequencing depth always enhances resolution, but the improvement quickly saturates. Beyond a sequencing depth of ten times the amount of DNA, much deeper sequencing is needed to significantly increase resolution. For comparison, we estimated the resolution for the nine datasets in Table 1 (Fig.4c). For each of them, we first estimated the noise extent by binning the sequences as we did before, to then estimate the initial DNA amount (Material and methods) and eventually the resolution. For some datasets, depending on the initial DNA amount, the sequencing depth was close to or beyond saturation, while for others, further improvement could still be possible.

To further investigate the saturation effect, we computed the exploited resolution (Materials and Method) by normalizing with respect to the different initial DNA amount, so as to focus on the distance from the saturation point (Fig.4d). Using the exploited resolution, we then defined an optimal sequencing depth *D*^opt^ as the depth at which the exploited resolution reaches 95%. This optimal depth can be computed as (Materials and methods)

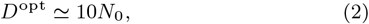

which means that the optimal NGS depth is about ten times larger than the amount of DNA before amplification (*N*_0_). The main message of our article is that, for a given initial DNA amount *N*_0_, increasing the sequencing depth beyond 10*N*_0_ offers no benefit due to the saturation of the exploited resolution. Direct experimental verification suggests that an even stricter rule could be considered (Fig. S6).

We computed the exploited resolution and the optimal depth *D*^opt^ for the nine datasets in Table 1. We observed that NGS depth was too large for three of the datasets, and too small for two of them (Fig. 4d). Using our rule of thumb (2), we can now recommend the appropriate NGS depth for measuring DNA barcode density based on the estimate total number of DNA barcodes for each scenario.

## Discussion

In this work we addressed the problem of determining an optimal NGS depth for sampling highly diverse libraries with barcodes. Optimal depth is usually discussed in terms of coverage, which sets the target reading redundancy for complete sequencing [21, 22, 23].

These established practices and methods are however designed for sequencing long DNA fragments, or even entire genomes. They may not be suitable for estimating the abundances of short read sequences, such as molecular barcodes, because of the overwhelming diversity of the sample. In these cases, NGS depth is often determined heuristically, or simply pushed at maximum, with additional costs and resources. Thanks to our analyses, we understood that increasing the sequencing depth above certain values does not increase the dataset quality. Based on our results, we found that the optimal depth is about ten times the initial DNA amount before amplification (*N*_0_, see Results).

Our “rule of thumb” for the sequencing depth does not depend on the properties of the barcode library, such as its diversity or flatness. This follows from the independence of the noise extent from those quantities: it is the same for all barcodes and depends only on the sequencing depth and the initial number of DNA molecules (Eq.1). As a consequence, the signal-to-noise ratio has the same dependence on the sequencing depth for each barcode, and - after a rescaling - this allows us to focus on resolution and its exploited percentage. These two quantities are therefore the same for all barcodes. Eventually, the saturating behaviour of the exploited resolution allowed us to derive our rule for optimal sequencing depth. Finally, although our rule of thumb does not depend directly on the number of unique barcodes, the latter still has an influence on it. Indeed, a highly heterogeneous library will require a larger number of initial barcodes to ensure sufficient coverage, which in turn necessitates a higher NGS depth according to our rule of thumb. In contrast, for highly biased libraries, a smaller number of initial barcodes is adequate, resulting in a lower NGS depth.

Increasing this DNA copy number always improves the resolution of an NGS dataset (Fig. 4c), yet for a given value, depths larger than the optimal value do not improve data quality (Fig. 4d), and in particular the ability to measure the concentration of DNA barcodes. For the application of our rule of thumb, the initial amount of DNA needs to be measured. Various methods are available for this purpose. One example is the NanoDrop, which enables rapid quantification of DNA molecules using only 1-2 µL of sample, with a relatively low error rate of 5% [47]. It is important to note that our rule of thumb is predicted on experimental conditions involving only two subsamplings: one prior to PCR amplifications and one subsequent to them (see Fig. 3a). Additional subsamplings may be necessary in certain experiments; in these cases, it is required to measure the concentration at each step (see Supplementary Mathematics for the derivation): *D*^opt^ ≃ 10*/*(Σ_*i*_ 1*/N*_*i*_). Based on this formula, one can derive recommended values for the NGS reads in any setting.

Our results come from a combination of data analysis of NGS datasets and mathematical modeling, which allowed us to quantify the over-dispersion in data (large noise extent), and to understand its nature. At first, for the NGS datasets of molecular barcodes (Table 1), we inferred the original concentrations of each barcode in the sample using a simple, yet precise model of the count probability distribution. This allowed us to study the count statistics in three datasets (Fig. 2), where variance and mean are proportional. The proportionality factor, identified as the noise extent of the dataset, is considerably higher (*AAV datasets*) or slightly higher (*hYAP65 WW dataset*) than 1 (Poisson noise). We then built a three-step mathematical model for NGS procedures, including the amplification process required to produce the necessary amount of genetic material [48]. We were able to solve the model analytically, obtaining a concise mathematical expression for the noise extent (Eq. 1). Our formula recovered the correct over-dispersion relation, and showed that reducing the number of amplification rounds has a limited impact on noise (Fig. 3e). From this we deduced that over-dispersion is an intrinsic property of NGS [48], that can at best be attenuated by accurate experimental protocols. Lastly, we observed that noise extent increases with sequencing depth. This is the reason for the small noise in the *hYAP65 WW datasets* and for the saturation of the resolution of count measurements (Fig. 4). From this last observation, we defined the optimal sequencing depth as the value at which the exploited resolution reaches 95 % of its saturated value, and subsequently we derived an analytical expression for that optimal value. What makes it possible to define a consistent sampling depth at the desired resolution is that mean and variance of the counts are proportional and that the proportionality factor is the same for all barcodes in the sample and it scales linearly with the depth.

Previous works in the literature already discussed and modeled noise in NGS data. They however limited their analysis on the statistical properties of noise, and described it with over-dispersed negative binomial [25, 26, 28, 29] or beta-binomial [27] probability distributions. Here we built a mathematical model to understand the origin of such over-dispersion. Our amplification model is based on a series of mathematical works studying PCR [31, 30, 32, 33, 34, 35], among which [32] was particularly relevant. We used their analytical frameworks to study the statistics of noise in NGS counts, thereby establishing a mathematical foundation for analyzing over-dispersion.

Three main limitations of our work can lead to further developments. First of all, the replication rate might be more complex than a Michaelis-Menten with constant parameter. However, this is a minor limitation, because it will not much impact the properties of the noise that we need to derive our optimal depth. Also, noise extent in NGS data showed a small dependence on the expected count. In our analyses, we observed small deviations for large count values (Fig. 2d and g). Going beyond our simple position independent model slightly reduced this effect (Fig. S2), and we expect that better models can reduce even more this deviation: errors in predicting the expected counts introduce some additional fluctuations in our binning procedure, and therefore increase the observed noise variance. In our analysis we did not try more flexible models because overfitting NGS counts lead to underestimation of noise extent, providing a trade off between model complexity and precision. It is also possible that different regimes appear for large count values [49] and the mechanism that generate these different regime remains unknown. In fact, previous work already noticed that the count variance grows faster than the mean [25, 26, 27, 28, 29]. Our mathematical analysis excluded these behaviors, but this might come from our simplified amplification process which considered only missed replications. Finally, other sources of errors might be present during PCR, such as those resulting in replacing one nucleotide for another during replication [48]. Even if these errors can be reduced by high-fidelity PCR and have in general smaller consequences in comparison to missed replication [24, 48], their integration into our framework is possible as a future perspective towards refinement of our method. Another interesting avenue to explore would be about optimizing the absolute signal-to-noise ratio and not the exploted one (which is mathematically equivalent to the exploited resolution). In this case, to estimate the predicted count of “rare” barcodes it would be necessary to increase the initial amount of DNA (*N*_0_), and the sequencing depth should be adjusted following our rule of thumb. We leave this last interesting point of discussion for future developments.

## Acknowledgments

The authors would like to thank L. C. Byrne for providing the AAV dataset 2. The authors would also like to thank J. Fernandez-de-Cossio-Diaz, G. Uguzzoni, L. C. Byrne and M. Desrosiers for useful comments and discussions. The authors would also like to thank Twist Bioscience for producing the plasmid library and the platform GENOM’IC at Institut Cochin for sequencing. This work was supported by ERC Starting Grant (REGE-NETHER 639888 to D.D.), European Research Council (ERC) Horizon 2020 Framework Programme Project (863214 – NEUROPA to D.D.), UNADEV, BpiFrance (Grant i-Demo - GEAR project to D.D. and U.F.), the Institut National de la Santé et de la Recherche Médicale (INSERM), Sorbonne Université (to D.D. and U.F.), The Foundation Fighting Blindness, Agence National de Recherche (ANR) RHU Light4-Deaf, LabEx LIFESENSES (ANR-10-LABX-65 to D.D.), IHU FOReSIGHT (ANR-18-IAHU-01 to D.D.), JSPS KAKENHI (Grant Number 22K17994 to T.N.), World Premier International Research Center Initiative (WPI), MEXT, Japan (to T.N.), and Paris Region Postdoctoral Fellowship (PRPF to E.Z.).

## Author contribution

Takahiro Nemoto and Ulisse Ferrari equally contributed.

## Data and code availability

The in-house dataset will be available upon acceptance. The code used for the analysis is available at this GitHub link: https://github.com/tommyocari/NGS

