## Supplementary figures and appendix for "Optimal sequencing depth for measuring the concentrations of molecular barcodes"

### Supplementary figure 1: the frequencies of nucleotides and of amino acids in the different datasets

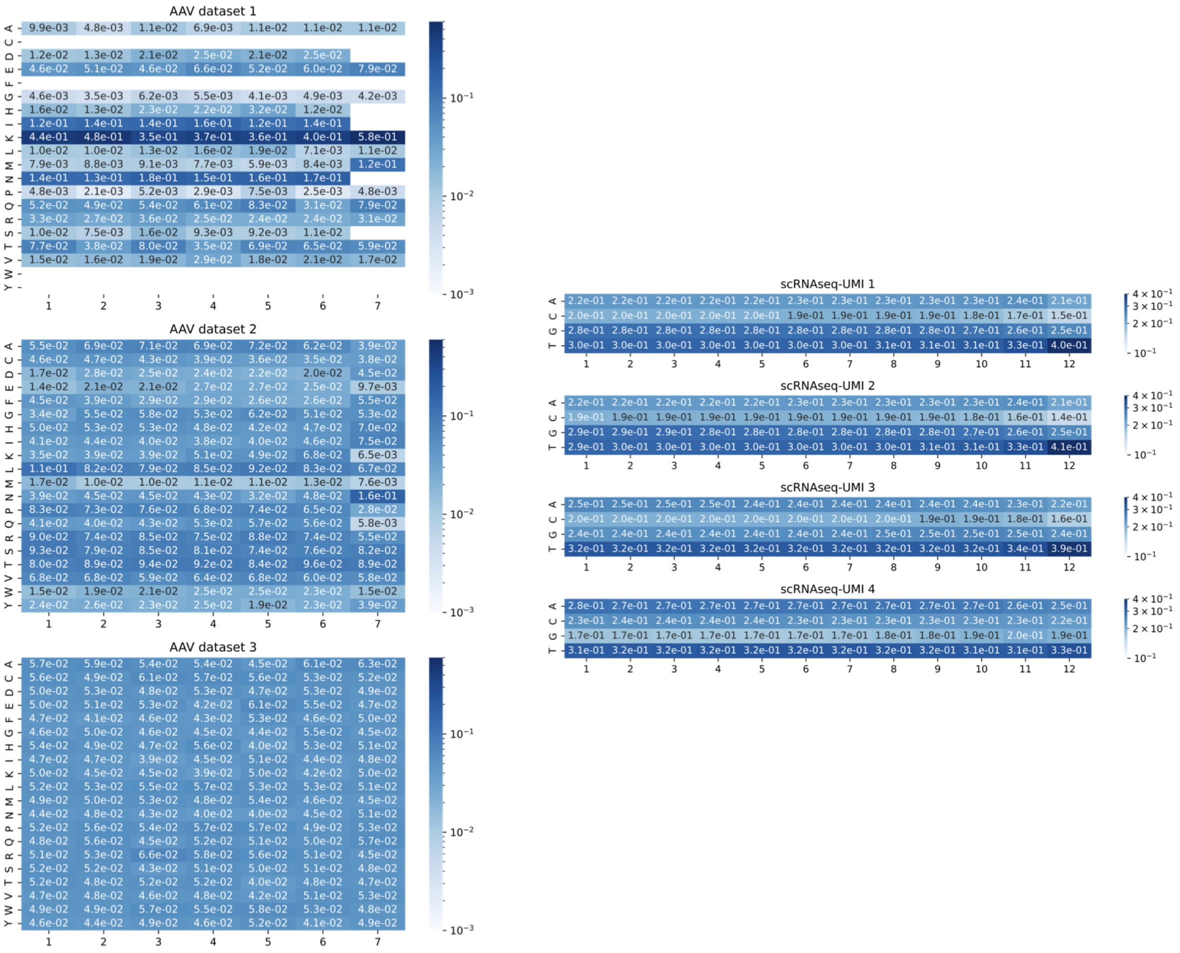

### Supplementary figure 2: parameters of the model for the error-prone PCR of the hYAP65 WW datasets and the analysis of the replicate 2

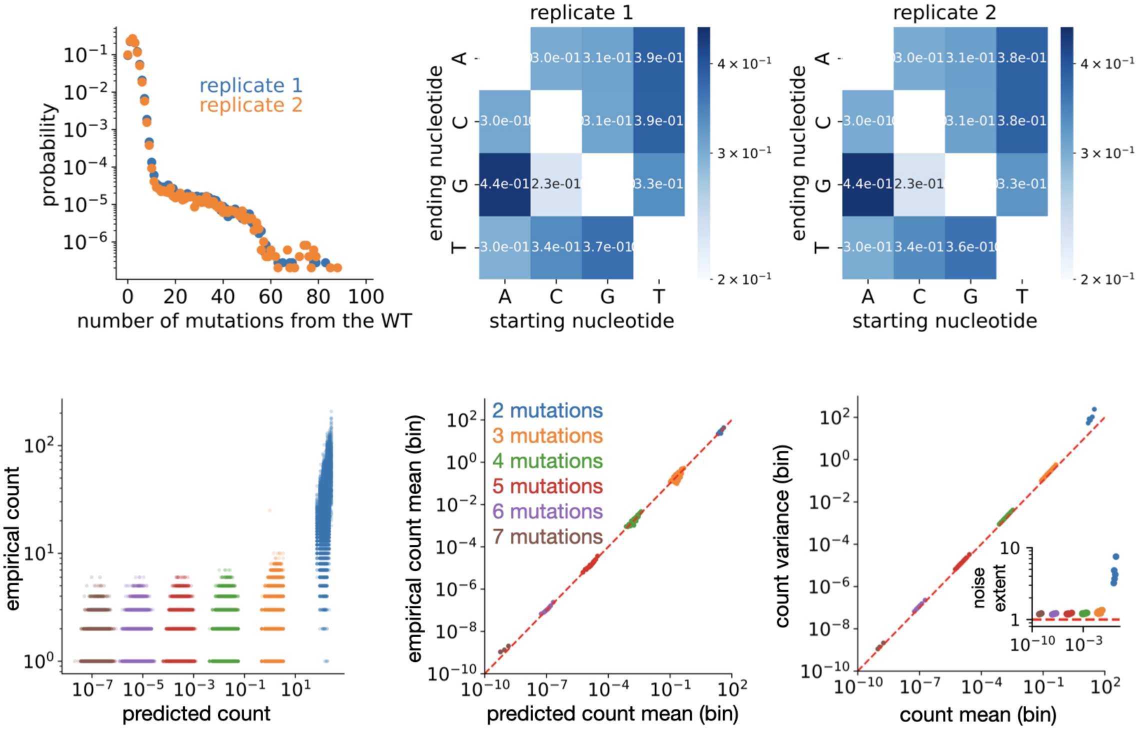


Top row: the parameters of the mutation model inferred on the two replicates of the hYAP65 WW dataset. The matrices shown represent the probability of mutation of a nucleotide in another nucleotide.

Bottom row: the same analysis performed in Fig. 2 on the dataset hYAP65 WW rep 2.

### Supplementary figure 3: the deviations from the proportionality law are due to the inexactitudes of the model

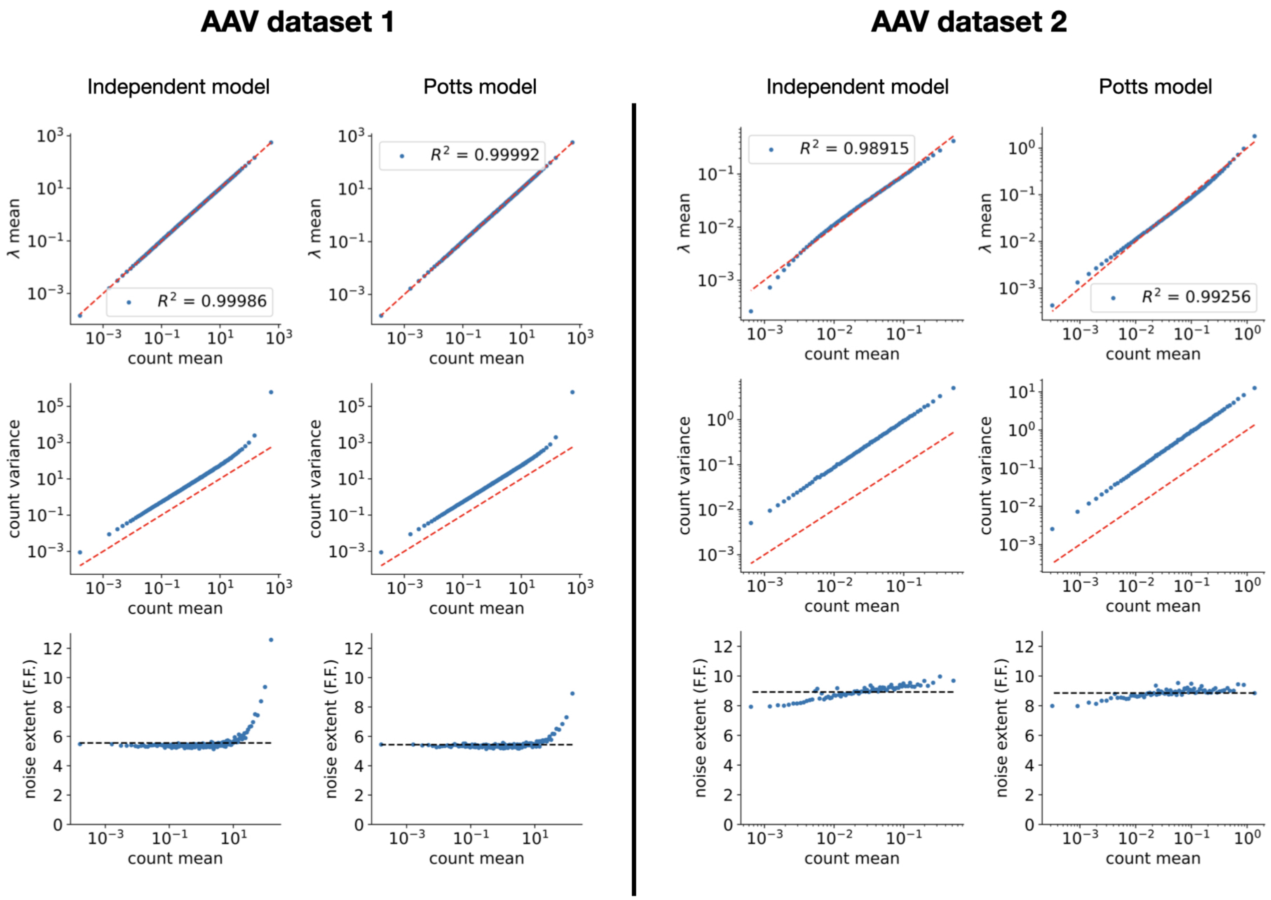


After inferring the position-independent model, we binned the data with respect to the expected count predicted by the model on the all possible sequences. In each bin, the sequences have similar expected count and each bin has an equal amount of sequences with nonzero empirical counts. We therefore plotted (see left column for both datasets) the mean of the expected count against the mean of the empirical count, the variance of the empirical counts against the mean of the empirical count and the noise extent (F.F.) computed as in Material and methods against the count mean. From the last plot we can see in both case slight deviations from the horizontal black dashed line (which represent the mean of the points). These deviations are due to the use of a simplified model. In fact we inferred a more complex model (like a Potts model inferred with the pseudo-likelihood methods), which is able to take into account the correlations between different positions. The same plots are done again with the other model. From the top panels (mean $\lambda$ vs mean count), we see that in both datasets Potts model is slightly outperforming the independent model. However the bottom panels shows that the improvement on the proportionality relationship is more marked.

### Supplementary figure 4: the mathematical framework analysed with constant duplication probability confirms the mean variance proportionality

$$p_n = p$$

$$\mathrm{Var}(c^i) = \left(1 + a\frac{D}{N_0}\right) E(c^i) \qquad \text{with} \qquad a = \frac{2}{1+p}$$

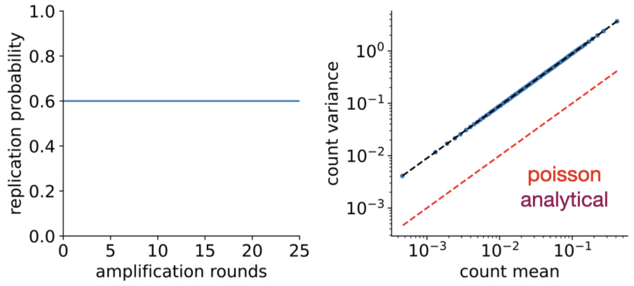


As a simplified case, the mathematical framework has been analysed in the case of constant duplication probability $p_n = p$. The mathematical computation recovered the proportionality relationship between mean and variance. The Fano factor (proportionality factor) has the same structure of the one obtained from the Michaelis Menten replication probability, with a more simple expression of $a = 2/(1+p)$. The dependencies on the NGS depth $D$ and the initial DNA molecules $N_0$ is the same as the ones obtained in the paper. We therefore ran a complete simulations of the whole mathematical framework with similar parameters as AAV dataset 2, generated a synthetic dataset and binned as explained in the first section of the Results. We plotted count variance against count mean for each bin (blue points). The simulated data are over dispersed, since they lay over the identity line (poisson noise, red dashed line) and they are in agreement with the analytical prediction (purple dashed line).

### Supplementary figure 5: the Fano factor dependency on the initial reduced DNA amount and on the amplification rounds

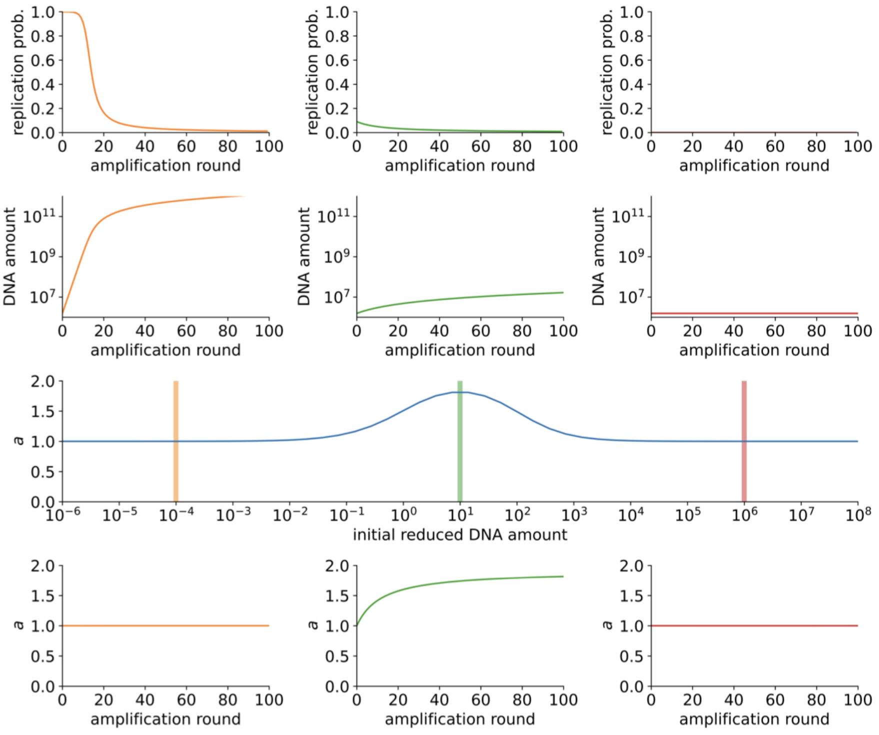


From the analytical calculation on the mathematical framework, we did not manage to get a simple analytical formula for the function $a$, so we wrote a simulation function which inputs the initial reduced DNA amount $N_0/K_M$ and outputs the probability of replication, the DNA polymerase ratio and the value of $a$ over 100 rounds of amplification. Thanks to this function it was possible to study the trend of $a$ with both the initial reduced DNA amount and the number of amplification rounds. Three regimes were found.

- The amplification regime (yellow) happens when $N_0/k < 10^{-1}$. The number of DNA molecules is strongly increasing and then reaches a linear regime. In this regime $a$ is not depending on the amplification rounds and it is equal to 1. This regime is the one where realistic PCR is.
- The under-amplification regime (green) happens when $10^{-1} < N_0/k < 10^3$. The number of DNA molecules is weakly increasing and the $a$ is increasing with the number of amplification. This means that the noise in this regime is high and increase with the number of amplification rounds, but this regime is not realistically happening since the amount of DNA increases very slowly during the PCR rounds.
- The non-amplification regime (red) happens when $N_0/k > 10^3$. Here the amplification of the DNA amount is linear and the $a$ is equal to 1 and not dependent on the number of amplification rounds.

#### Supplementary figure 6: direct experimental validation of our rule of thumb

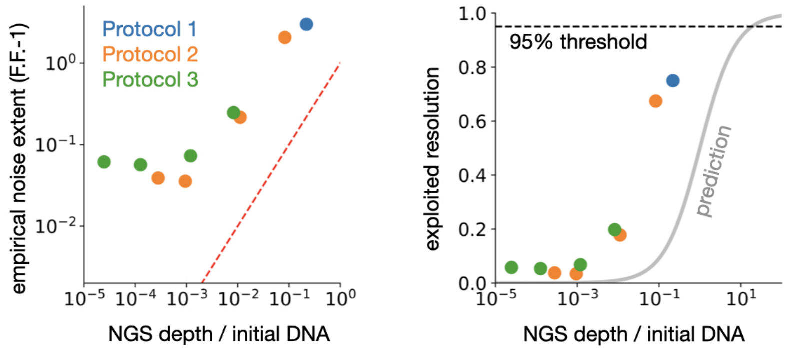


Fig. S6. Left: empirical noise extent plotted against NGS sequencing depth divided by initial DNA amount, for different experimental preparations of the same biological sample (AAV dataset 3). Right: empirical (colored points) and model-predicted (gray line) exploited resolution plotted with respect to NGS sequencing depth divided by initial DNA amount.

In this analysis, we aimed to experimentlally validate our noise model. To do this, we performed a set of experiments in which both the sequencing depth (D) and the number of initial DNA barcodes before amplification ($N_0$) are varied. More specifically, we used our in-house dataset (AAV dataset 3), composed of AAV2 plasmids, and prepared four diluted samples (100 ng, 10 ng, 1 ng, 0.1 ng), corresponding respectively to $N_0 = 10^{10}, 10^9, 10^8, 10^7$. These samples were then sent to two different NGS facilities and sequenced based on the following three different protocols:

- **Protocol 1**: high-depth sequencing (D~240M reads, $N_0 \sim 10^9$) at Facility 1, referred to as *AAV dataset 3* in the previous figures of the paper.
- **Protocol 2**: shallow sequencing (D~100k reads, $N_0 \sim 10^{10}, 10^9, 10^8, 10^7$) at Facility 2.
- **Protocol 3**: medium-depth sequencing (D~1M reads, $N_0 \sim 10^{10}, 10^9, 10^8, 10^7$) at Facility 2.

We started by analysing how the noise extent in the data depends on sequencing depth and initial DNA amount, and showed that deeper sequencing leads to greater noise extent (left panel), thereby validating one of our main results. Additionally, note that the ratio of sequencing depth to initial DNA amount ($D/N_0$) corresponds to our model's prediction of the noise extent itself (red line in the figure and Eq. (1) in the paper). As expected, the empirical noise is larger than our predictions, likely due to additional sources of noise that are not accounted for in our model (see discussion). The right panel shows the exploited resolution, computed from the empirical data (see below), as a function of the ratio $D/N_0$. It also includes a comparison between the empirical results (colored points) and model predictions (grey line). Note that the empirical points and the predicted line differ by an offset, which corresponds to the larger variability observed in the data (left panel). As a consequence, our rule of thumb - “don't sequence beyond 10 times the initial DNA amount” - is supported, although the data suggest that an even stricter rule could be possible. Additionally, when plotted against $D/N_0$, the empirical exploited resolutions (colored points) collapse onto a single curve, confirming that this ratio is the key quantity to consider.
Empirical exploited resolution is computed by rewriting its formula (see methods) in terms of noise extent (F.F.), and then using the value estimated from the data (y-axis of left panel).
Overall, these results show how our model captures the empirical behavior of the sequencing noise, and support its use for deriving our rule of thumb.

#### Supplementary mathematics

##### Analytical derivation of the mean-variance proportionality

Let's start by following the replication of one DNA variant along $n$ amplification cycles. We start by a large heterogeneous initial library from which a number of DNA fragments are taken as the initial sample for PCR. The initial copy number of variant $i$, $C_0^i$, is a random variable with mean $E_0^i = \mathbb{E}\left(C_0^i\right)$ and variance $V_0^i = Var\left(C_0^i\right)$. Let's suppose this random variable is Poissonian distributed, so that $V_0^i = E_0^i$. At each iteration, each copy of variant $i$ are replicated with probability $p_n$, which in principle depends on the amplification round, therefore their number grows as:

$$C_n^i = C_{n-1}^i + B[C_{n-1}^i, p_n] \ , \tag{1}$$

where $B[C, p]$ is a binomial random variable with parameters $C$ and $p$. On average we have:

$$E_n^i = (1 + p_n)E_{n-1}^i, \tag{2}$$

Computing the variance is slightly harder:

$$\begin{aligned} V_n^i &= \text{Var}\left(C_{n-1}^i + B[C_{n-1}^i, p_n]\right) = \\ &= \text{Var}\left(C_{n-1}^i\right) + \text{Var}\left(B[C_{n-1}^i, p_n]\right) + \\ &\quad +2\text{Cov}(C_{n-1}^i, B[C_{n-1}^i, p_n]) = \\ &= V_{n-1}^i + p_n(1-p_n)E_{n-1}^i + p_n^2 V_{n-1}^i + 2p_n V_{n-1}^i = \\ &= p_n(1-p_n)E_{n-1}^i + (1+p_n)^2 V_{n-1}^i \end{aligned} \tag{3}$$

##### Case where $p_n$ is independent of $n$

In the case where $p_n = p$ does not depend on the amplification round, the equation (2) gives:

$$E_n^i = (1+p)^n E_0^i \tag{4}$$

Deplacing the (4) in the (3) and calling $q = 1 - p$ and $s = 1 + p$ we find:

$$\begin{aligned} V_n^i &= s^2 V_{n-1}^i + pqs^{n-1}E_0^i \\ V_{n-1}^i &= s^2 V_{n-2}^i + pqs^{n-2}E_0^i \cdot \left(s^2\right)^1 \\ V_{n-2}^i &= s^2 V_{n-3}^i + pqs^{n-3}E_0^i \cdot \left(s^2\right)^2 \\ &\cdots\cdots \quad \cdots\cdots \\ V_2^i &= \quad s^2 V_1^i + pqs^1 E_0^i \quad \cdot \left(s^2\right)^{n-2} \\ V_1^i &= \quad s^2 V_0^i + pqs^0 E_0^i \quad \cdot \left(s^2\right)^{n-1} \end{aligned}$$

Summing the all the left-hand side and the right-hand side many terms are simplifying, resulting in:

$$\begin{aligned} V_n^i &= \left(s^2\right)^n V_0^i + pqE_0^i \left(\sum_{k=0}^{n-1} \left(s^2\right)^k s^{n-1-k}\right) \\ &= s^{2n} V_0^i + pqs^{n-1}E_0^i \left(\sum_{k=0}^{n-1} s^k\right) \\ &= \left[s^n + \frac{pq}{s}\left(\sum_{k=0}^{n-1} s^k\right)\right] s^n E_0^i \\ &= \left[s^n + \frac{pq\left(s^n - 1\right)}{s(s-1)}\right] E_n^i \end{aligned}$$

where in the third line $V_0^i = E_0^i$ is used.

Now, for $n$ big enough $s^n \gg 1$, so:

$$V_n^i = \left(1 + \frac{q}{s}\right) s^n E_n^i$$

After the amplification, a portion of the DNA strands is NGS sequenced for a total of $D$ reads. Therefore, the measured count $c^i$ of variant $i$ will be Poissonian distribute with parameters:

$$c^i \sim \text{Poisson}\left(\frac{DC_n^i}{\sum_j C_n^j}\right) \simeq \text{Poisson}\left(\frac{DC_n^i}{s^n \sum_j E_0^j}\right)$$

where we discarded the fluctuations of the denominator thanks to the central limit theorem.

From the above equation and after calling $N_0$ the total number of DNA strands at the beginning of the amplification process, it follows:

$$\begin{aligned} \mathbb{E}\left(c^i\right) &= \frac{D}{s^n N_0}\mathbb{E}\left(C_n^i\right) = \frac{D}{N_0}E_0^i \\ \text{Var}\left(c^i\right) &= \mathbb{E}\left(c^i\right) + \text{Var}\left(\frac{DC_n^i}{(1+p)^n \sum_j E_0^j}\right) \\ &= \mathbb{E}\left(c^i\right) + \frac{D^2}{s^{2n}N_0^2}\text{Var}\left(C_n^i\right) \\ &= \mathbb{E}\left(c^i\right) + \frac{D^2}{s^{2n}N_0^2}\left(1 + \frac{q}{s}\right) s^n E_n^i \\ &= \mathbb{E}\left(c^i\right) + \frac{D}{N_0}\left(1 + \frac{q}{s}\right)\mathbb{E}\left(c^i\right) \end{aligned}$$

By replacing back the values of $q$ and $s$ we find the equation:

$$\text{Var}\left(c^i\right) = \left(1 + \frac{2}{1+p}\frac{D}{N_0}\right) \cdot \mathbb{E}\left(c^i\right)$$

This equation, once replaced $a = 2/(1+p)$, is the one shown in the Results of the paper.

##### Case where $p_n$ is dependent of $n$

In the more general case $p_n$ does depend on the amplification round, the equation (2) gives:

$$E_n^i = \prod_{k=1}^{n} \left(1 + p_k\right) E_0^i \tag{5}$$

Replacing the (5) in the (3) and calling $q_n = 1 - p_n$, $s_n = 1 + p_n$ and $\mathscr{S}_n = \prod_{k=1}^{n}\left(1 + p_k\right)$ we find:

$$
\begin{aligned}
V_n^i =& \quad s_n^2 V_{n-1}^i + p_n q_n \mathscr{S}_{n-1} E_0^i \\
V_{n-1}^i =& s_{n-1}^2 V_{n-2}^i + p_{n-1} q_{n-1} \mathscr{S}_{n-2} E_0^i \cdot \left(s_n^2\right) \\
V_{n-2}^i =& s_{n-2}^2 V_{n-3}^i + p_{n-2} q_{n-2} \mathscr{S}_{n-3} E_0^i \cdot \left(s_n^2 s_{n-1}^2\right) \\
\ldots\ldots & \qquad \ldots\ldots \\
V_2^i =& \quad s_2^2 V_1^i + p_2 q_2 \mathscr{S}_1 E_0^i \qquad \cdot \left(s_n^2 s_{n-1}^2 \ldots s_3^2\right) \\
V_1^i =& \quad s_1^2 V_0^i + p_1 q_1 \mathscr{S}_0 E_0^i \qquad \cdot \left(s_n^2 s_{n-1}^2 \ldots s_3^2 s_2^2\right)
\end{aligned}
$$

Summing the all the left-hand side and the right-hand side many terms are simplifying, resulting in:

$$
\begin{aligned}
V_n^i =& \left(\mathscr{S}_n\right)^2 V_0^i + E_0^i \sum_{k=0}^{n-1} \mathscr{S}_k p_{k+1} q_{k+1} \prod_{m=k+2}^{n} s_m^2 \\
=& \left(\mathscr{S}_n\right)^2 V_0^i + E_0^i \sum_{k=0}^{n-1} p_{k+1} q_{k+1} \prod_{m=1}^{k} s_m \prod_{m=k+2}^{n} s_m \prod_{m=k+2}^{n} s_m \\
=& \left(\mathscr{S}_n\right)^2 V_0^i + E_0^i \sum_{k=0}^{n-1} \frac{p_{k+1} q_{k+1}}{s_{k+1}} \mathscr{S}_n \prod_{m=k+2}^{n} s_m \\
=& \left(\mathscr{S}_n\right)^2 V_0^i + E_0^i \mathscr{S}_n \sum_{k=0}^{n-1} \frac{p_{k+1} q_{k+1}}{s_{k+1}} \prod_{m=k+2}^{n} s_m \\
=& \left[\mathscr{S}_n + \sum_{k=0}^{n-1} \frac{p_{k+1} q_{k+1}}{s_{k+1}} \prod_{m=k+2}^{n} s_m\right] \mathscr{S}_n E_0^i \\
=& \left[\mathscr{S}_n + \sum_{k=0}^{n-1} \frac{p_{k+1} q_{k+1}}{s_{k+1}} \prod_{m=k+2}^{n} s_m\right] E_n^i \\
=& w_n E_n^i
\end{aligned}
$$

where in the fifth line $V_0^i = E_0^i$ is used and in the last line we call $w_n$ the expression in parenthesis.

After the amplification, a portion of the DNA strands is NGS sequenced for a total of $D$ reads. Therefore, the measured count $c^i$ of variant $i$ will be Poissonian distribute with parameters:

$$
c^i \sim \text{Poisson}\left(\frac{D C_n^i}{\sum_j C_n^j}\right) \simeq \text{Poisson}\left(\frac{D C_n^i}{\mathscr{S}_n \sum_j E_0^j}\right)
$$

where we discarded the fluctuations of the denominator thanks to the central limit theorem. From the above equation and after calling $N_0$ the total number of DNA strands at the beginning of the amplification process, it follows:

$$
\begin{aligned}
\mathbb{E}\left(c^i\right) =& \frac{D}{\mathscr{S}_n N_0} \mathbb{E}\left(C_n^i\right) = \frac{D}{N_0} E_0^i \\
\text{Var}\left(c^i\right) =& \mathbb{E}\left(c^i\right) + \text{Var}\left(\frac{D C_n^i}{(1+p)^n \sum_j E_0^j}\right) \\
=& \mathbb{E}\left(c^i\right) + \frac{D^2}{\left(\mathscr{S}_n\right)^2 N_0^2} \text{Var}\left(C_n^i\right) \\
=& \mathbb{E}\left(c^i\right) + \frac{D^2}{\left(\mathscr{S}_n\right)^2 N_0^2} w_n E_n^i \\
=& \mathbb{E}\left(c^i\right) + \frac{D}{N_0} \frac{w_n}{\mathscr{S}_n} \mathbb{E}\left(c^i\right)
\end{aligned}
$$

By replacing $a = w_n / \mathscr{S}_n$ we find the following equation:

$$
\text{Var}\left(c^i\right) = \left(1 + a \frac{D}{N_0}\right) \cdot \mathbb{E}\left(c^i\right)
$$

This equation is the one shown in the Results of the paper.

#### Extension: multiple subsampling

Most of the biological experiments are made by more than two pipetting subsampling. Let's suppose at each of those subsamplings we pipette out a DNA amount of respectively $N_0, N_1, N_2, \ldots, N_{M-1}$. Since, as result of the paper, the role of the amplification steps is negligible at first order, we can consider only multiple iterative Poisson sampling. We are going to find the general formula in a recursive way. Let's denote with $c_n^i$ the count of the $i$-th sequence coming from the $n$-th Poisson sampling:

$$
c_n^i \sim \text{Poisson}\left(\frac{c_{n-1}^i}{N_{n-1}} N_n\right)
$$

The average of the random variable is equal to:

$$
\mathbb{E}\left(c_n^i\right) = \frac{N_n}{N_{n-1}} \mathbb{E}\left(c_{n-1}^i\right)
$$

The variance is obtained via the theorem of the total variance:

$$
\begin{aligned}
\text{Var}\left(c_n^i\right) &= \text{Var}\left(\frac{c_{n-1}^i}{N_{n-1}} N_n\right) + \mathbb{E}\left(\frac{c_{n-1}^i}{N_{n-1}} N_n\right) \\
&= \left(\frac{N_n}{N_{n-1}}\right)^2 \text{Var}\left(c_{n-1}^i\right) + \frac{N_n}{N_{n-1}} \mathbb{E}\left(c_{n-1}^i\right)
\end{aligned}
$$

We know that for $M = 2$ total Poisson samplings, the variance is proportional to the mean, so let's suppose that for each $n$ and $i$, $\text{Var}\left(c_n^i\right) = F_n \cdot \mathbb{E}\left(c_n^i\right)$ and let's replace it in the equation:

$$
\begin{aligned}
\text{Var}\left(c_n^i\right) &= \left(\frac{N_n}{N_{n-1}}\right)^2 F_{n-1} \cdot \mathbb{E}\left(c_{n-1}^i\right) + \frac{N_n}{N_{n-1}} \mathbb{E}\left(c_{n-1}^i\right) \\
&= \frac{N_n}{N_{n-1}} F_{n-1} \cdot \mathbb{E}\left(c_n^i\right) + \mathbb{E}\left(c_n^i\right)
\end{aligned}
$$

We can rewrite it as an iterative equation for the proportionality factor:

$$
F_n = 1 + \frac{N_n}{N_{n-1}} F_{n-1}
$$

The solution of this iterative formula, after calling $D = N_{M-1}$, is:

$$
F_n = 1 + D \cdot \sum_m \frac{1}{N_m}
$$
